# Cerebellar climbing fibers convey perceptual choice during decision-making

**DOI:** 10.1101/2025.02.06.636959

**Authors:** Tin Long Yiu, Evan A. Wilson, Takaki Yahiro, Lei Ma, Maozhen Qin, Haining Zhong

## Abstract

Cerebellar climbing fibers are thought to signal reward prediction errors in non-motor functions. By imaging postsynaptic responses of climbing fibers onto mouse Purkinje cell dendrites during auditory discrimination, we found that climbing fibers in crus I can encode cue identities or perceptual choices. These responses were reshaped by reversal learning. Optogenetic perturbation of climbing fiber activity impaired discrimination. These results suggest a feedforward role of climbing fibers in perceptual decision-making.

---

The cerebellum is traditionally associated with motor control. Recent evidence from human and primate studies suggests that it also plays a critical role in cognitive functions^1–3^. However, the types of cognitive information being processed in the cerebellum and the pathways involved remain not fully understood. Recent work has shown that climbing fibers (CFs), which constitute one of the two main inputs to the cerebellum and instruct plasticity changes in Purkinje cells (PCs)^4,5^, carry feedback reward prediction errors (RPEs)^6^ in several cerebellar lobules^7,8^. Nevertheless, the climbing fiber activity in lobule crus I does not conform to the framework of either RPE or motor-related signaling^9^. It is also unclear how these activities evolve during cognitive learning. Here, we examine the climbing fiber activity of crus I during an auditory cognitive task in which mice learn to associate similar sensory stimuli with different reward outcomes. Our results indicate that a fraction of climbing fibers carries feedforward signals encoding information of subjective decision, which is essential for the execution of learned cognitive behaviors.

To explore the cognitive function of climbing fibers, we developed an auditory association task for head-fixed mice compatible with two-photon imaging and optogenetics (Fig. 1a). A trace period was added between the cue period and reward outcome to isolate potential neuronal activities related to decision-making from those associated with reward retrieval. In the initial “generalization” stage, water-deprived mice learned to associate both low- (8 kHz) and high-frequency (14 kHz) tones with water reward. After training (5 sessions), mice exhibited similar lick responses to both cues (licking in > 90% of trials; Fig.1b-d; Extended Data Fig. 1). They were then advanced to the “discrimination” stage where only the 8-kHz tone was paired with reward. Mice rapidly learned to discriminate between the tones by decreasing their licking response to the 14-kHz tone to less than 30% of trials over 3–7 sessions (Fig.1b-d; Extended Data Fig. 1). This paradigm enabled us to assess climbing fiber responses during context-dependent perceptual decisions evoked by comparable auditory stimuli.

**Fig. 1.**
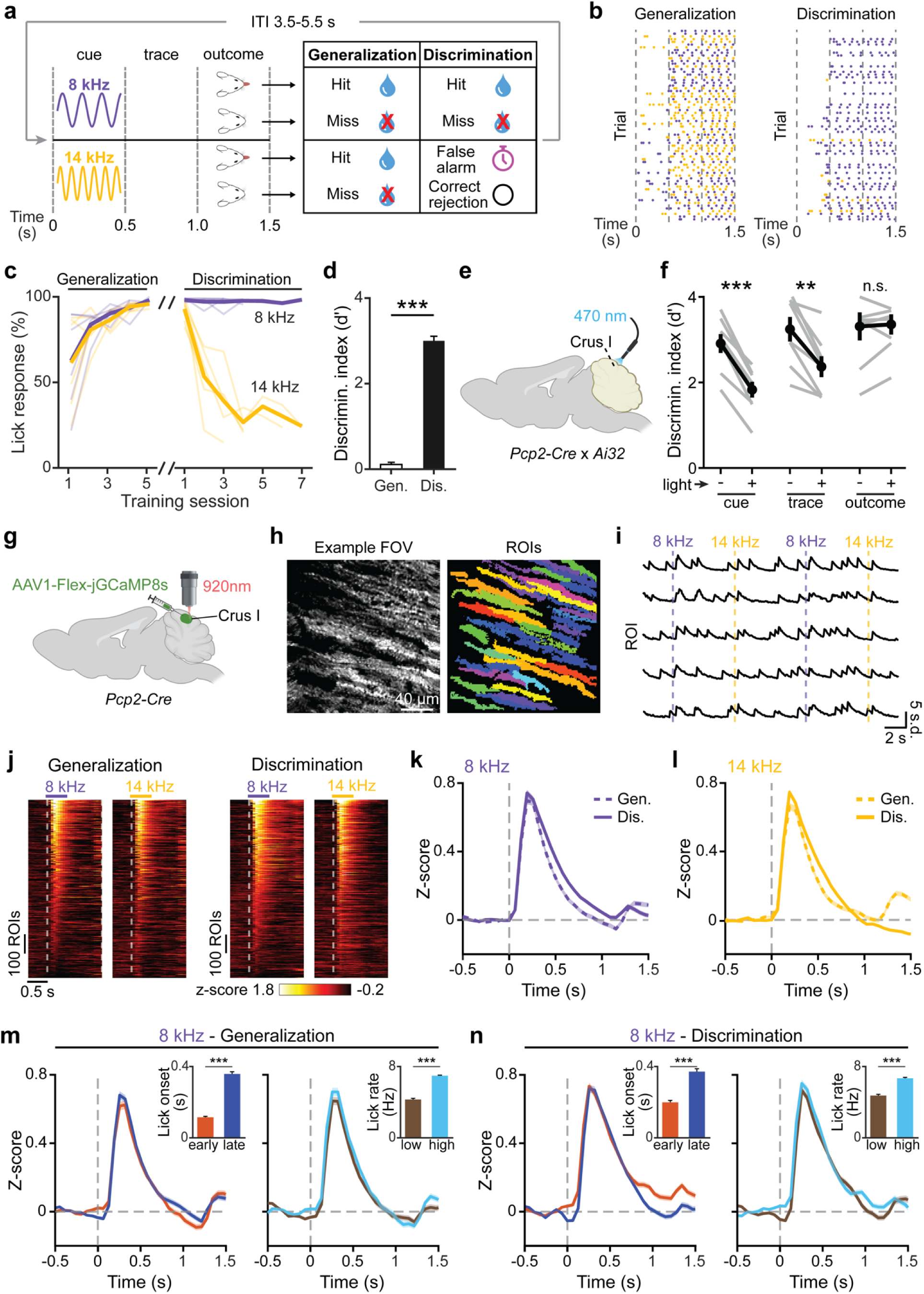
Enhanced cue-evoked climbing fiber responses during sensory discrimination. **a**, Schematic representation of the two-stage auditory association task. **b**, Example lick responses for individual sessions during generalization and discrimination. Each dot represents a lick. **c**, Learning curve of lick responses to each cue across training sessions of generalization and discrimination. **d**, Behavioral performance (quantified as discrimination index d’) for all imaging sessions during generalization and discrimination. The same set of fields of view (FOVs) were examined (*n* = 31 from 6 mice; Two sample t-test; *P* = 4.14 × 10^−33^). **e**, Schematic of optogenetic stimulation of Crus I Purkinje cells in *Pcp2-Ai32* mice. **f**, Discrimination performance with or without PC stimulation during cue, trace, and outcome period (*n* = 8 mice, paired t-test, *P* = 2.61× 10^−5^, *P* = 0.0016, and *P* = 0.73, respectively). **g**, Schematic illustrating AAV1-Flex-jGCaMP8s virus injection into crus I and two-photon imaging setup. **h**, Representative image of jGCaMP8s expression in Purkinje cell dendrites in an example FOV (left) and extraction of dendritic ROIs (right). **i**, Example calcium activity traces for 8 kHz and 14 kHz trials from individual ROIs. **j**, Raster plots showing averaged calcium activity in response to 8 kHz and 14 kHz tones in all dendritic ROIs during generalization and discrimination. *n* (ROIs/mice) = 1010/6 (generalization) and 1056/6 (discrimination). **k**, **l**, Baseline-subtracted average calcium responses during 8 kHz (**k**) and 14 kHz (**l)** trials across stages. **m**, **n**, Baseline-subtracted average calcium responses stratified by lick onset (earliest 25% vs. latest 25%; left) and lick rate (highest 25% vs. lowest 25%; right) in 8 kHz trials during generalization (**m**) and discrimination (**n**). Insets display average lick onset and average lick rate, respectively. From left to right, *P* = 2.88 × 10^−53^, 4.14 × 10^−62^, *P* = 4.98 × 10^−^17, and *P* = 1.38 × 10^−^25, Two samples t-test. From left to right, *n* = 249, 280, 253, 284, 278, 309, 280, and 311 trials. All error bars and error bands represent the s.e.m. and their centers represent the mean. n.s.: *P* > 0.05; \*\**P* ≤ 0.01; \*\*\**P* ≤ 0.001.

We focused on lobule crus I, which receives auditory inputs^10,11^ and its climbing fiber activities do not follow RPE characteristics in a conditioning behavior^9^. To test crus I’s involvement in the task, we optogenetically stimulated channelrhodopsin-expressing Purkinje cells in trained *Pcp2-Cre; Ai32* double heterozygous mice during the discrimination stage (Fig. 1e). Photostimulation (470 nm, 1.2 mW/mm^2^, 0.5 s) of crus I during the cue and trace periods, but not the outcome periods, significantly impaired discrimination performance (Fig. 1f), which was primarily due to increased licking to the non-rewarding 14-kHz tone (Extended Data Fig. 2a–d). Importantly, PC stimulation alone minimally triggered licking and off-window stimulation did not result in perturbation in discrimination (Extended Data Fig. 2e, f), suggesting that crus I is involved in cognitive processing during the task.

To monitor climbing fiber activity, we imaged calcium dynamics in Purkinje cell apical dendrites, which faithfully reflect individual climbing fiber inputs^12,13^. Expression of jGCaMP8s was achieved in the left crus I of *Pcp2-Cre* mice by injecting a Cre-dependent adeno-associated virus (AAVs). The fluorescence of jGCaMP8s was imaged *in vivo* using a two-photon microscope in the Purkinje dendrites during the behavioral task and extracted using Suite2p^14^ from regions of interest (ROIs) (Fig. 1g–i). During both generalization and discrimination, the majority of ROIs (*n* = 738/1010 and 874/1056, respectively) exhibited increased CF inputs in response to both tones (Fig. 1j). Across conditions, the calcium response peaked during the cue period (0-0.5 s) and gradually decayed over the trace period. A smaller calcium response was observed during the outcome period except for the 14 kHz tone during the discrimination stage, likely reflecting reward retrieval (Fig. 1k, l; Extended Data Fig. 3). These responses during the cue and trace period were independent of lick onset or vigor, indicating that they were not driven by licking movement (Fig. 1m, n). Notably, average calcium activities for both tones during the cue and trace periods were similar, or slightly increased, at the discrimination stage (Fig. 1k, l; Extended Data Fig. 3b-c). These results confirm that climbing fiber responses in crus I are not RPEs^6,9^, which would have diminished for the non-rewarded 14 kHz tone during the cue and trace periods.

To understand the task variables contributing to climbing fiber activities, we fit a generalized linear model (GLM) to the averaged calcium response of individual ROIs during the cue and trace periods (0–1 s after cue onset) by using tone identity and lick count as the predictors (Methods). Following discrimination learning, GLM weights for tone identity increased while those for lick decreased (Fig. 2a, b). Furthermore, the GLM weights for tone identity were consistently larger than those for lick, and the difference increased from generalization to discrimination (Fig. 2c). These results suggest that a major component of the climbing fiber activity is driven by tone discrimination.

**Fig. 2.**
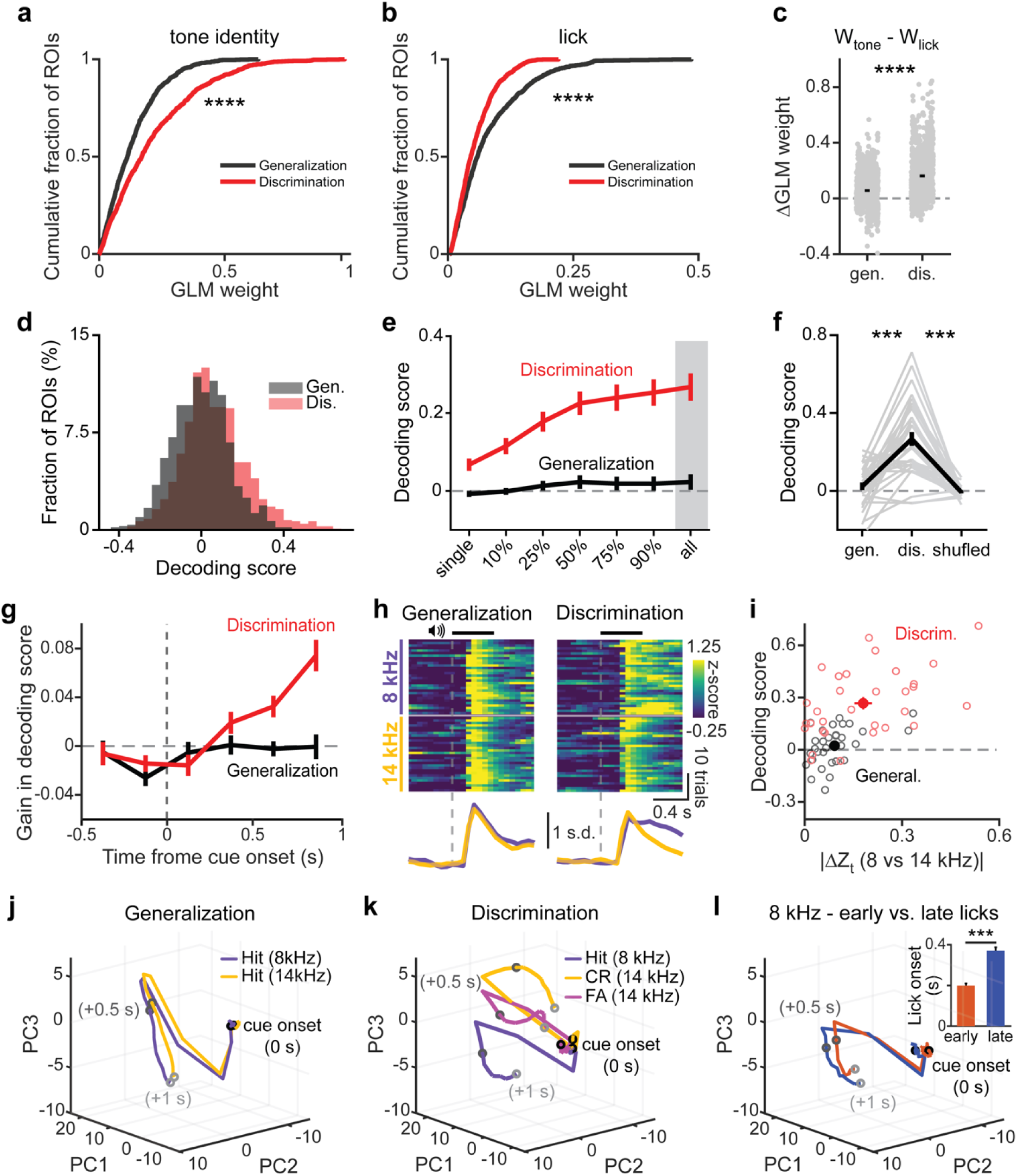
Populational climbing fiber inputs encode decision-related information. **a,** Cumulative distribution of tone identity GLM weights across stages (Kolmogorov–Smirnov test, *P* = 5.43 × 10^−20^). **b,** Cumulative distribution of lick GLM weights across stages (Kolmogorov– Smirnov test, *P* = 5.14 × 10^−13^). **c,** Difference between tone identity and lick GLM weights across stages (Two sample t-test, *P* = 6.40 × 10^−56^). **d**, Decoding score distribution for individual dendritic ROIs across stages (Kolmogorov–Smirnov test, *P* = 1.31 × 10^−12^). **e**, Decoding scores averaged across FOVs showing improved performance of SVM with increasing population size (from 10% to all ROIs) only during discrimination. **f**, Changes in decoding scores for FOVs across stages, with tone labels shuffled as controls (Paired t-test*, P* = 5.10 × 10^−7^ and 1.25 × 10^−8^, respectively). **g**, Temporal contributions of climbing fiber inputs to gain in decoding scores during generalization and discrimination. **h**, Raster plots (top) of single-trial populational climbing fiber activity in a representative FOV and the trial-averaged traces (bottom) during generalization (left) and discrimination (right). **i**, Correlation between decoding scores and absolute differences in climbing fiber inputs for 8 and 14 kHz tones across generalization and discrimination. **j**, Population neural activity trajectories for correct 8 kHz and 14 kHz trials during generalization. **k**, Population neural activity trajectories for hit, correct rejection (CR), and false alarm (FA) trials during discrimination. **l**, Population neural activity trajectories for the earliest 25% and latest 25% licking 8 kHz trials during discrimination. Average lick onsets are shown in inset (*n* = 278 and 309, respectively; Two sample t-test*, P* = 4.98 × 10^−17^). All error bars and error bands represent the s.e.m. and their centers represent the mean. \*\*\**P* ≤ 0.001. *n* (ROIs/mice) = 1010/6 (generalization) and 1056/6 (discrimination).

To dissect the spatiotemporal encoding of tone information, we trained a support vector machine (SVM)^15,16^ to classify tone identity on a trial-by-trial basis using the corresponding climbing fiber activities from individual ROIs (Methods). During generalization, the average decoding score across ROIs was near zero (i.e., at chance level) (Fig. 2d, e). However, discrimination learning shifted the overall scores higher (Fig. 2d). Although the average gain for individual ROIs was modest, the inclusion of increasing number of ROIs per FOV significantly improved the decoding during discrimination but not during generalization or when the tone identity was shuffled (Fig. 2e, f). This suggests that either tone or decision information is encoded at the populational level. Alignment of FOVs across animals in crus I showed no apparent spatial organization of the decoding capability (Extended Data Fig. 4). To examine the temporal contribution, we systematically dropped data from individual time bins (Methods). Time bins closer to the end of the trace period contributed increasingly to the decoding accuracy (Fig. 2g). To understand how the information may be encoded in calcium activity, which on average was only subtly different between generalization and discrimination, we averaged the calcium activity of all neurons within individual FOVs during a given trial. We found prominent calcium activity differences during the trace period in a fraction of FOVs (Fig. 2h). Although these differences can be positive or negative, their absolute magnitude correlated with the decoding capability of the respective FOV (Fig. 2i).

The above results indicate that the climbing fiber activity distinguishes between different tone stimuli. However, this distinction could reflect either the tone identity or the animal’s perceptual decision to lick or not. To disentangle these possibilities, we pooled ROIs from all FOVs and performed principal component analysis (PCA) on concatenated average activity traces from correct 8-kHz or 14-kHz trials (i.e., no miss or false alarm trials). We focused on the first three principal components, which explained over 80% of the data variance (Extended Data Fig. 5a, b). In this PCA space, trajectories for correct 8-kHz and 14-kHz trials at the generalization stage were similar (Fig. 2j; Extended Data Fig. 5c, f). However, during discrimination, the trajectories diverged ∼300 ms after cue onset (Fig. 2k; Extended Data Fig. 5d, f). This divergence could not be attributed to licking movements because the trajectories of 8-kHz trials with the earliest and latest quartiles of lick initiation times were nearly identical (Fig. 2l; Extended Data Fig. 6d, e), as were those with the highest and lowest quartiles of lick rates (Extended Data Fig. 6a–c). Notably, the trajectory for false-alarm 14-kHz trials also diverged at a similar time, but it fell between the hit and correct rejection traces (Fig. 2k; Extended Data Fig. 5e, f). These results may be explained if the climbing fibers carry information about both the perceptual choice and sensory cue such that, in false-alarm trials, the perceptual choice (to lick) may deviate from the actual cue sensation (14 kHz) and result in a trajectory that reflects a blend of the two.

To dissect how the information is encoded in individual climbing fibers, we calculated the difference in calcium signals during the trace period between 8 and 14 kHz trials and correlated these differences with the corresponding decoding score (Fig. 3a, b). Compared to generalization, discrimination learning greatly increased the number of ROIs that exhibited significant differences, either positive or negative, beyond an arbitrary threshold of 2x s.d. of the data at generalization (23% vs 6%, *P* < 0.001). These discriminative ROIs achieved significantly higher decoding scores in the SVM model (Fig. 3c). To understand what drives the differences, we applied the MLSpike algorithm^17^ to convert fluorescence traces of individual trials into climbing fiber activation events (Extended Data Fig. 7a). Consistent with earlier populational level results (Fig. 2g, h), the major difference between generalization and discrimination phases arose from changes in the likelihood of climbing fiber activation events occurring from late cue period to the trace period (Fig. 3d, e).

**Fig. 3.**
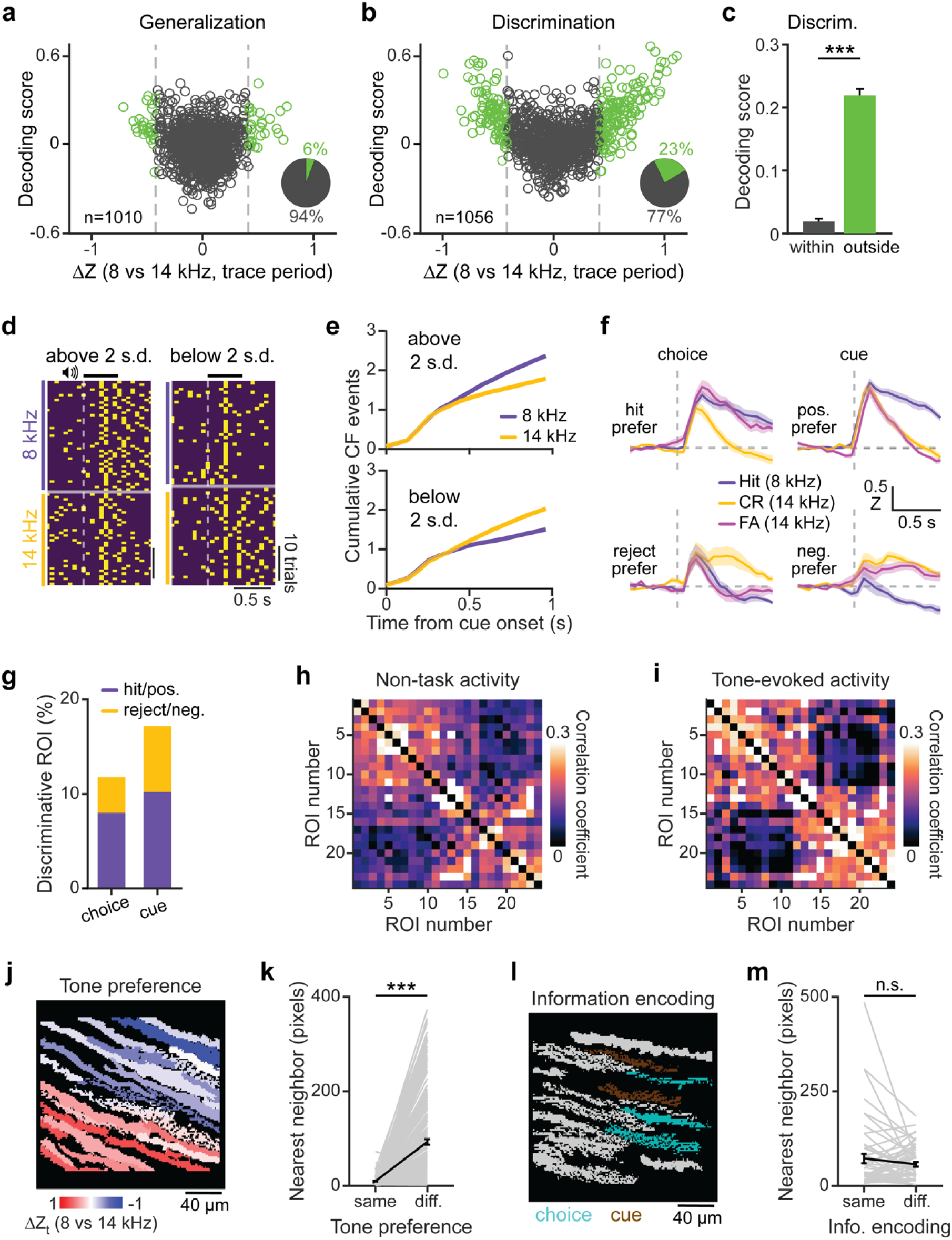
Climbing fiber inputs in individual Purkinje cell dendrites encode choice or cue identity. **a-b**, decoding score and tone-response differences for dendritic ROIs during generalization (**a**) and discrimination (**b**). Discriminative ROIs (beyond ±2 SD from the mean difference during generalization) are highlighted in green. Insets display the fraction of discriminative ROIs. **c**, Mean decoding scores for discriminative ROIs (outside 2 s.d.; *n* = 246) and non-discriminative ROIs (within 2 s.d; *n* = 810) during discrimination (Two sample t-test; *P* = 2.42 × 10^−78^). **d**, Raster plots of single-trial climbing fiber activation events for an 8 kHz preferred (left) and a 14 kHz preferred ROI (right). **e**, Cumulative climbing fiber activation for all 8 kHz preferred ROIs (top) and 14 kHz preferred ROIs (bottom). **f**, Trial-averaged traces for ROIs encoding choice (left) and cue (right) with hit/8 kHz preference (top) and correct rejection/14 kHz preference (bottom). **g**, Fraction of discriminative ROIs encoding choice or cue. Colors indicate hit/8 kHz preference or correct rejection/14 kHz preference. **h, i**, Correlation matrices of fluorescence activity during non-task (h) and tone-evoked periods (i) for an example FOV with ROIs sorted by location.. **j**, Spatial distribution of the tone-response differences in a representative FOV. **k**, Nearest-neighbor distances for ROIs with the same or different tone preference (*n* = 221 ROI, paired t-test, *P* = 5.61 × 10^−31^). **l**, Spatial distribution for ROIs encoding choice or cue identity in a representative FOV. **m**, Nearest-neighbor distances for ROIs with the same or different information encoding property (*n* = 54 ROI, paired t-test, *P* = 0.2170). All error bars and error bands represent the s.e.m. and their centers represent the mean. n.s.: *P* > 0.05; \**P* ≤ 0.05; \*\*\**P* ≤ 0.001. |

To dissect how choice and cue identities are differentially encoded, we compared the calcium responses in false alarm trials (i.e., the animal licked for the non-rewarding 14-kHz tone) with correct hit or rejection trials in the discriminative ROIs. We found that ∼29% of these ROIs exhibited a “choice” encoding pattern in which the false alarm response mirrored correct hit responses or a “cue” encoding pattern in which the false alarm responses resembled the correct rejection responses (Fig. 3f, g; Extended Data Fig. 7b–d; Methods).

We next asked how these ROIs were spatially organized. The cerebellum is known to feature zonal and microzonal organizations^18^ (see also Extended Data Fig. 7e). We performed a correlation analysis of calcium activity within individual FOVs and found that the microzonal boundaries were evident even when using only background activities outside of behavioral trials (i.e., within ITI; Fig. 3h, i). Dendrites with similar tone preferences appeared to be spatially clustered such that their nearest neighbor of the same response type was much closer than dendrites with opposite responses (Fig. 3j, k). Surprisingly, dendrites encoding choice or cue were intermingled (Fig 3l, m), revealing that there is functional heterogeneity within individual microzones. Overall, our results indicate that a subset of climbing fibers in crus I encode information related to the subjective decision to act, while some others encode the identity of auditory cues.

To consolidate the above conclusion and to determine whether climbing fiber activities may change during learning, we reversed the cue-reward association in a subset of mice following the discrimination phase by instead pairing the 14-kHz tone, but not the 8-kHz tone, with reward (Fig 4a; Extended Data Fig. 8). Consistent with our earlier results, the 8-kHz tone continued to evoke climbing fiber responses even in the absence of reward association (Extended Data Fig. 8d), further supporting that these responses do not represent RPEs. Interestingly, principal component analysis of the reversal data revealed that activity trajectories for correct 8-kHz and 14-kHz trials also diverged but in the opposite direction (Fig 4b; Extended Data Fig. 8g-j) compared to the discrimination stage (c.f. Fig. 2k). SVM analysis showed that the decoding scores of individual ROIs shifted towards negative when tone labels supplied to the model were kept the same as in the prior stage (Fig. 4c; Method). At the FOV level, the populational activity of dendritic ROIs could decode the tone identity (Fig. 4d; Extended Data Fig. 8e–f), and the decoding score correlated with the absolute difference in calcium responses between the two tones (Fig. 4e). To examine plasticity in individual climbing fibers, we tracked specific ROIs across generalization, discrimination, and reversal stages. Approximately 24% of ROIs were visually matched (Methods) across conditions (Fig. 4f). We found that overall many ROIs in the discrimination phase switched their response preference upon cue-reward reversal (Fig.4g–i; Extended Data Fig. 9a). Thus, climbing fiber activity can adapt to changes in task contingencies and flexibly encodes decision-related information.

**Fig. 4.**
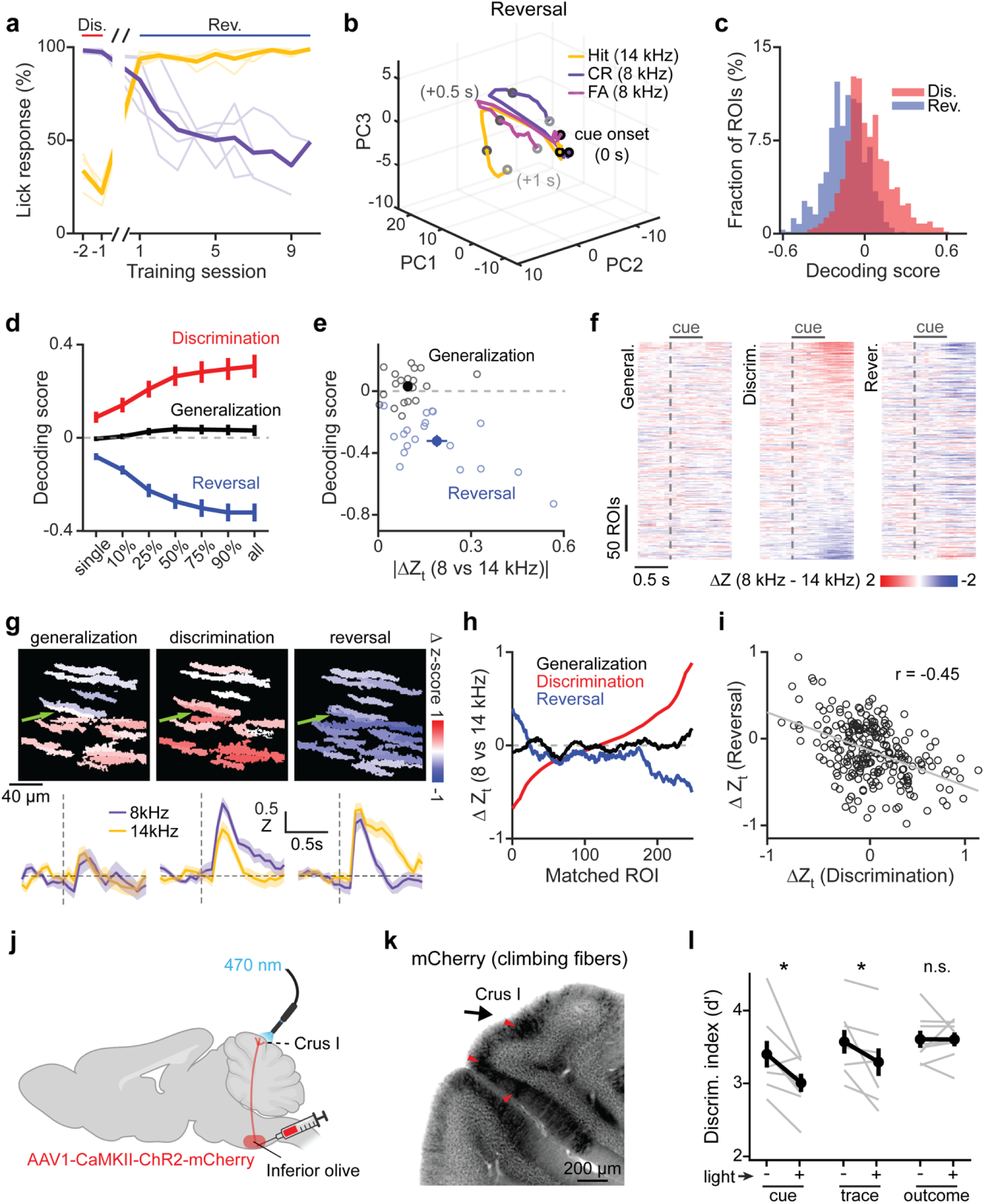
Plasticity and causal role of climbing fibers in perceptual decision-making. **a**, Learning curve of lick responses to each cue across the last two training sessions of discrimination and all sessions of reversal. **b**, Population neural activity trajectories of hit, correct rejection (CR), and false alarm (FA) trials during reversal (*n* = 646 ROIs). **c**, Decoding score distribution for individual dendritic ROIs across stages (*n* = 617 for discrimination and *n* = 646 for reversal; Kolmogorov–Smirnov test, *P* = 3.62 × 10^−42^). **d**, Averaged decoding scores across FOVs for different population sizes in the SVM. **e**, absolute differences in climbing fiber inputs for 8 and 14 kHz tones across generalization and reversal. **f**, Raster plots of calcium response differences for all matched dendritic ROIs (*n* = 247) during generalization (left), discrimination (middle), and reversal (right). **g**, Calcium response differences for matched ROIs in an example FOV (top) across stages and the trial-averaged traces (bottom) of an example polarity-switching ROI (green arrow). **h**, Mean calcium response differences during trace period for matched ROIs across stages. **i**, Correlation of calcium response differences for matched ROIs between discrimination and reversal (Pearson’s Linear Correlation Coefficient, *P* = 1.45 × 10^−13^). **j**, Schematic illustrating AAV1-CaMKII-ChR2-mCherry virus injection into inferior olive and optogenetic stimulation of Crus I climbing fiber axons. **k**, Example mCherry expressions in climbing fibers (arrows). **l**, Discrimination performance with or without climbing fiber stimulation during cue, trace, and outcome period (*n* = 8 mice, paired t-test, *P* = 0.021, *P* = 0.040, and *P* = 0.99, respectively). All error bars and error bands represent the s.e.m. and their centers represent the mean. n.s.: *P* > 0.05; \**P* ≤ 0.05; \*\*\**P* ≤ 0.001.

Finally, we asked whether proper climbing fiber activity in crus I was essential for decision execution. Since opto-stimulation of Purkinje cells (Fig. 1e, f) might perturb both simple and complex spikes^19^, we injected ChR2-mCherry under a CaMKIIα promotor in the inferior olive and optogenetically stimulated climbing fibers in crus I during the discrimination task (Fig 4j, k). Photostimulation during cue and trace periods significantly impaired discrimination performance (Fig. 4l). This effect was primarily due to an increase in the false alarm rate among 14 kHz trials and could not be attributed to light perception or motor artifacts (Extended Data Fig. 10), suggesting that climbing fiber inputs into the cerebellum is essential in decision execution.

Taken together, our results reveal that a fraction of crus I climbing fibers conveys information about the sensory cue identity or the perceptual choice of action. This information is acquired during learning and is essential for accurate decision execution. These conclusions expand our understanding of the functional role of climbing fibers, which are thought to encode feedback errors, i.e. RPEs, in the context of cognitive function^7–9^. In contrast, a perceptual choice suggests an unappreciated, feedforward role of climbing fibers in cognitive decision. This notion is strengthened by the observation that the decision-related information is acquired during learning and is stable once learned (Fig. 3a–h; Extended Fig. 7d). The information is expressed before the outcome occurs (Fig. 2g, k; 3e, f). Since no moment-by-moment feedback is involved, our results indicate that, in contrast to other cerebellar regions where climbing fibers provide RPE-like signals^9^, crus I may operate during decision-making in a mode described by the inverse internal model^20,21^, in which the cerebellar circuit learns from feedforward cortical commands and sensory inputs to eventually be able to estimate the cortical response under a given situation. This functional difference might stem from distinct connectivity patterns across cerebellar lobules^18^. Crus I is highly interconnected with the prefrontal cortex^22^, which could provide cognitive command signals to the climbing fiber pathway via the mesodiencephalic junction.

In addition, our findings suggest that climbing fibers with similar tone-response preferences are spatially clustered resembling microzones^18,23^ (Fig. 3h–k). However, dendritic ROIs encoding choice-related or cue-related information are intermingled (Fig. 3l, m), revealing that functional heterogeneity may exist within individual microzones. Furthermore, our findings highlight the adaptability of climbing fiber encoding. At the reversal stage, climbing fibers flexibly adjusted to new cue-reward associations primarily by switching their response polarity (Fig. 4a–h). This flexible encoding likely supports the cerebellum’s capacity to adapt to continuously changing demands. Finally, while climbing fiber inputs are known to instruct cerebellar plasticity^4,5^, our findings suggest that these activities themselves can also undergo learning-dependent changes. Overall, our results suggest a feedforward role of climbing fibers in cognitive processing.

## Methods

### Experimental animals

All animal experiments and handling were conducted in accordance with the US National Institutes of Health Guide for the Care and Use of Laboratory Animals and were approved by the Institutional Animal Care and Use Committee (IACUC) of the Oregon Health & Science University (#IP00000420).

Two-photon imaging experiments were performed on *Pcp2(L7)-Cre* mice (*n* = 6), maintained in-house by crossing to *C57/BL6J* wild-type mice. Optogenetics stimulation of Purkinje cells experiments were performed using *Pcp2(L7)*-*Ai32* mice (i.e., *Pcp2(L7)-Cre* mice crossed with *Ai32* reporter line; *n* = 8). Optogenetic stimulation of climbing fibers experiments were performed using *C57/BL6J* wild-type mice (*n* = 8 mice). Both female and male mice were included the study, all aged >P60. Mice were group housed on a 12:12 light-dark cycle, with male mice occasionally single-housed after surgery to prevent fighting.

### Surgery

Mice were intraperitoneally injected with dexamethasone (5 mg/kg) and buprenorphine (1 mg/kg) before surgery to minimize brain swelling and postoperative pain. Mice were then anesthetized with isoflurane (4% for induction and 1-1.5% for maintenance) and mounted on a stereotaxic frame with body temperature regulated with a heating pad. A custom-made head plate was affixed over the left cerebellar folium Crus I (coordinates relative to Lambda: antero-posterior, −3.5 mm; medio-lateral, +1.5 mm) and secured with dental cement (Metabond, Parkell). A 3 mm craniotomy was then performed at the center of the head plate to expose the cerebellar cortex. For imaging experiments, a Cre-dependent GCaMP8s virus (pGP-AAV-CAG-FLEX-jGCaMP8s-WPRE) diluted at 1:20 from stock titer was injected at 3 locations within Crus I. At each site, 300 nL of virus solution was pressure-injected at depths of 200-300 μm below the cerebellar surface. To prevent viral reflux, the injection pipette was held in place for 10 minutes before retraction. A 3 mm single-paned coverslip was press-fit into the craniotomy, sealed to the skull using cyanoacrylate (VetBond), and fixed in place by dental cement. For optogenetic stimulation of climbing fibers, an AAV expressing CaMKII-ChR2 (AAV1-CaMKIIa-hChR2(H134R)-mCherry) was pressure-injected into the right inferior olive (coordinates relative to Bregma: antero-posterior, −6.5 mm; medio-lateral, +0.4 mm; dorso-ventral, +5.6 mm). No virus was injected for optogenetic stimulation of Purkinje cells experiments. To minimize light scattering during optogenetic stimulation, black pigment was incorporated into the dental cement, and non-Crus I regions were covered with a layer of black dental cement. Additionally, the conical portion of a nitrile rubber seal (RS Components) was then glued to the head plate with dental cement to block mice from directly viewing the LED light during photostimulation. The cranial window was filled with Kwik-Cast for protection during recovery and between experiment sessions.

### Behavioral training

Following a minimum recovery period of 7 days post-surgery, mice were placed on water restriction for 2 days and maintained at ∼80–85% of their initial body weight throughout the experiments. Typically, mice first underwent 2 days of habituation sessions where each lick on a waterspout was rewarded with 3 µL of saccharin solution (1.4mg/L), delivered at a maximum rate of 2 µL/s. Once mice reliably initiated licking, they were trained to perform an auditory generalization task. In each trial of the generalization task, either an 8 kHz or 14 kHz auditory tone (75 dB, 0.5 s duration) was presented. Mice were rewarded with 3 µL of saccharin solution for licking during an outcome window 1–1.5 s after tone onset (Hit1 and Hit2 for 8 kHz and 14 kHz trials, respectively). Upon achieving an expert-level performance (Hit rate >80% for both trial types), mice progressed to an auditory discrimination task. During the discrimination task, 8 kHz tones were rewarded with 3 µL of saccharin solution for licking during the outcome window (Hit). Conversely, licking in response to 14 kHz tones during the outcome window resulted in a 4 s timeout punishment (false alarm). No reward or punishment was given for a lack of licking in response to 8 kHz (miss) or 14 kHz tones (correct rejection). Trials were pseudo-randomized with a 1:1 ratio of 8 kHz to 14 kHz tones, with a maximum of three consecutive trials with the same tone. Performance of mice was assessed using the sensitivity index (d’). The d’ value was calculated as

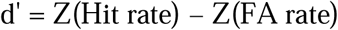

where Z represents the inverse of the cumulative distribution function of a normalized distribution whose mean is 0 and standard deviation is 1, as calculated using the *norminv* function in MATLAB. To avoid infinite d′ values, a Hit rate of 100% in a training session was adjusted to 99%. Mice were trained until their d’ exceeded 2.0. A subset of mice (*n* = 4) was further trained to perform a reversal task, where the reward contingency of the two tones was flipped such that licking in response to 8 kHz resulted in reward whereas licking in response to 14 kHz resulted in timeout punishment. Behavioral tasks were coordinated by Bpod (Sanworks) and custom MATLAB codes.

### Optogenetic experiments

*Pcp2(L7)-Ai32* mice (*n* = 8) and wild-type mice injected with CaMKII-ChR2 virus (*n* = 8) were trained to achieve a performance level of d’ >2 during the discrimination stage prior to optogenetic experiments. During the experimental session, each mouse performed 300 trials with photostimulation applied in 30% of trials (for both 8 and 14 kHz). Photostimulation was delivered using a 470 nm LED coupled with an optical fiber (M470F3, Thorlabs) directed at the cranial window at an intensity of 1.2 mW/mm^2^. Photostimulation was applied during distinct task epochs (cue, trace, or outcome) in separate sessions, with the duration of photostimulation matching the specific epochs length (500 ms). In control experiments, photostimulation was applied in 20% of trials without tone presentation (“LED only”) or with the optical fiber positioned outside the cranial window (“off window”).

### Behavioral analysis

Lick rate (Extended Data Fig. 1a, b; Extended Data Fig. 2a-c; Extended Data Fig. 8c; Extended Data Fig. 10a-c) was calculated as the average number of licks within a 50 ms time bin across trials. Lick response (Figure 1c; Extended Data Fig. 2d; Extended Data Fig. 10d) was quantified as the fraction of trials with at least one lick during the outcome window (1.0-1.5 s after cue onset). Lick onset was determined as the time between cue onset to the first lick in a trial (Figure 1m, n; Extended Data Fig. 6d, e).

### Two-photon imaging

Two-photon imaging was performed using a custom-built two-photon microscope based on the open-source MIMMS design from Howard Hughes Medical Institute Janelia Research Campus. The microscope was equipped with a 16X/0.8 NA objective lens (Nikon) and controlled using ScanImage software. Two-photon excitation was achieved with an 80 MHz pulsed Titanium-Sapphire laser at a wavelength of 920 nm. The emission was filtered using a Chroma ET500/40m-2p barrier filter and recorded with a cooled GaAsP photomultiplier tube (Hamamatsu H10769PA-40). Laser power at the sample was adjusted to be less than 40 mW. A 190 x 190 mm field of view was scanned using a galvanometer at a resolution of 512 x 128 pixels, with 8000 frames acquired at 15.625 frames/s during each imaging session. Imaging was performed across multiple FOVs spanning the medial-lateral axis of the left Crus I.

### Imaging analysis

Imaging data were processed using MATLAB (R2023b, MathWorks). Motion artifacts in the x-y plane were corrected, and dendritic regions of interest (ROIs) were extracted using Suite2P^14^. ROIs corresponding to PC dendrites were manually selected using the following criteria: 1. ROI shape excluded small, round, or vessel-like regions. 2. Mean fluorescence of the ROI exhibited characteristic calcium transients (rapid rise and slow decay) indicative of climbing fiber inputs. 3. Fluorescence signals avoided saturation, such as sustained fluorescence increases lasting several seconds. For the generalization and discrimination stages, we extracted 32 ± 7 and 34 ± 9 ROIs per FOV, respectively (mean ± SD, *n* = 31 FOVs from 6 mice; Fig. 1j). Calcium traces were z-scored, and correlation matrices were computed using the Pearson correlation coefficient between fluorescence traces of medial-lateral-aligned ROIs during non-task periods (outside 0–1.5 s from cue onsets; Fig. 3h) or the cue and trace period (Fig. 3i). Distances between ROIs were computed as perpendicular distances between a linear regression line fitted to the centroid of ROI based on the average dendritic angle of the FOV. ROIs were visually tracked across generalization, discrimination, and reversal stages. This was carried out by two independent experts and only mutually agreed ROI matches were included. The two tracking results were compared, and we only included ROIs that were identified as matches by both experts (∼24%) in the analysis (Fig. 4f-i; Extended Data Fig. 9).

### Generalized linear model (GLM) analysis

We modeled average climbing fiber inputs (0–1 s from cue onset) for individual dendritic ROIs as a function of tone identity (8 or 14 kHz) and lick count in the trial as in the following equation:

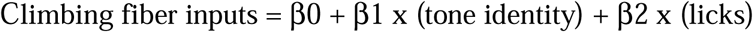

where β represents the coefficients. The MATLAB function fitglm was used, and we assume a normal distribution for climbing fiber inputs. GLM weights were defined as the absolute value of the fitted coefficients (β).

### Support vector machine (SVM) analysis

We train an SVM model for binary classification of tone identity (8 or 14 kHz) in each trial based on the respective climbing fiber inputs (-0.5 s to 1.0 s from cue onset) using the MATLAB function *fitcsvm*. We randomly split 80% of the trial for training the SVM model and evaluated the model performance using the remaining 20% of trials over 100 iterations. For single dendritic ROI analysis (Figure 2d; Figure 4c), input data comprised a row vector of climbing fiber input across time bin. For populational level analysis (Figure 2e; Figure 4d), input data was a matrix with rows representing dendritic ROIs, and columns representing time bins. Training data subsets (10%, 25%, 50%, 75%, 90%, or 100% of rows) were selected randomly. Model performance was validated by shuffling tone identity labels. Temporal contributions were assessed by systematically removing 6 sets of time bins (relative to cue onset: -0.512 s to -0.32 s, -0.256 s to -0.064 s, 0 s to 0.192 s, 0.256 s to 0.448 s, and 0.512 s to 0.704 s, 0.768 s to 0.96 s) and retraining the model. Decoding scores ranged from 0 (chance, 50% accuracy) to 1 (perfect accuracy) or -1 (perfect decoding with flipped tone identity in reversal stages).

### Neural activity trajectory analysis

To visualize neural trajectories of different trial outcomes, we first computed and concatenated the trial-averaged trace of correct 8 and 14 kHz trials such that each ROI’s calcium traces (-0.5 s to 1.0 s from cue onset) was represented as a 2T x 1 vector, where T is the total number of time bins per trial outcome type. We then combined all ROIs into a single matrix with dimensions 2T x N where N is the total number of ROIs. We used PCA for dimensionality reduction and projected the data onto a 3D neural subspace using the coefficient of the first three principal components and visualize trajectories of Hit1/Hit2 trials for generalization and Hit/CR/FA trials for discrimination and reversal. Variance explained by PCA components was confirmed to exceed chance by shuffling time bins (Extended Data Fig. 5a, b; Extended Data Fig. 8k). For lick onset and lick rate analysis (Figure 2l; Extended Data Fig. 6), we projected the data to 8 kHz trials with the earliest 25% and the latest 25% lick onset or the highest 25% and lowest 25% trials lick rate. For similarity analysis (Extended Data Fig. 5f; Extended Data Fig. 6c, e), we normalized the projected data such that the sum of all time-bins equal to 1 and computed the dot product between vectors of different trial types, where values of 1, 0, and -1 indicated identical, orthogonal, or opposite directions, respectively.

### Detection of climbing fiber activation events

We used an event detection algorithm, MLspike^17^, to identify fast dendritic calcium transients, faithful indicators of climbing fiber activation events in Purkinje cells (Extended Data Fig. 7). We used ΔF plus the maximum value of each ΔF trace (ΔF_max_) as an input to MLspike. Raw fluorescence traces were transformed to ΔF using the following equation: (F – F0), where F represents the raw fluorescence signal, F0 is the 8th percentile of fluorescence values within a 1-second window centered on each time point (±8 frames, with a total of 16 frames at 64 ms per frame). The baseline fluorescence parameter (F0) was set to ΔF_max_, the sampling rate (dt) was set at 1/15.625, and the indicator decay parameter (tau) was set to 0.15. The estimated height of a single climbing fiber activation event detected in each trace was set to 0.65. The output of MLspike is an event time. To prevent multiple counting, events detected in consecutive bins were consolidated by assigning them to the first time point of each sequence.

### Classification of discriminative ROI and its information encoding property

To identify discriminative ROIs, we calculated calcium signal differences during the trace period for 8 and 14 kHz tone stimuli across both generalization and discrimination stages. ROIs exhibiting signal differences exceeding ±2 standard deviations of the population mean during the generalization stage were classified as discriminative. For each discriminative ROI, baseline-subtracted calcium traces (0–1 s from cue onset) were calculated and grouped into three matrices corresponding to Hit, CR, and FA trials, with dimensions of Trial × Time. Pairwise two-sample t-tests were performed across time bins for each comparison (Hit vs. CR, Hit vs. FA, CR vs. FA). A time bin was considered significant if *P* < 0.05 for at least two consecutive bins.

Discriminative ROIs were further classified based on their response profiles: ROIs with a profile of “1-0-1” (significant for Hit vs. CR and Hit vs. FA but not CR vs. FA) were defined as choice-encoding; ROIs with a profile of “1-1-0” (significant for Hit vs. CR and Hit vs. FA but not CR vs. FA) were defined as cue-encoding. These classifications were used to analyze the encoding properties of climbing fiber inputs during generalization, discrimination, and reversal stages (Extended Data Fig. 7b).

## Acknowledgements

We thank the entire Zhong and Mao labs for discussions and helps, Drs. Stephen David, Skyler Jackman, Tianyi Mao and Tatsuo Sato for helpful comments on the manuscript. This work was supported by two NIH BRAIN Initiative awards R01NS104944 and RF1MH130784 (H.Z.), a NINDS R01 grant R01NS127013 (H.Z.), a Ronni Lacroute fellowship (T.L.Y.), a N.L. Tartar Trust fellowship (T.L.Y.) and a Sigma Xi Grants in Aid of Research (T.L.Y.).

## Author Contributions

TLY and HZ and conceived the project. TLY designed the experiments with HZ’s inputs. TLY carried out most experiments. EAW performed the AAV injection into the inferior olive. TY verified ROI matching. LM contributed to histology. MZ maintained and bred mice. TLY analyzed the data with HZ’s inputs. TLY and HZ wrote the manuscript, with edits and comments by all authors. HZ secured the funding and supervised the project.

## Data Availability

Numerical source data for plots will be provided with this paper. Original raw data will be provided upon request, in order to include all supporting information.

## Code Availability

Custom MATLAB codes will be made available upon request.

## Competing Interests

The authors declare no competing interests.

## Additional information

Correspondence and requests for materials should be addressed to H.Z.

## Extended Data Figures and Legends

**Extended Data Fig. 1.**
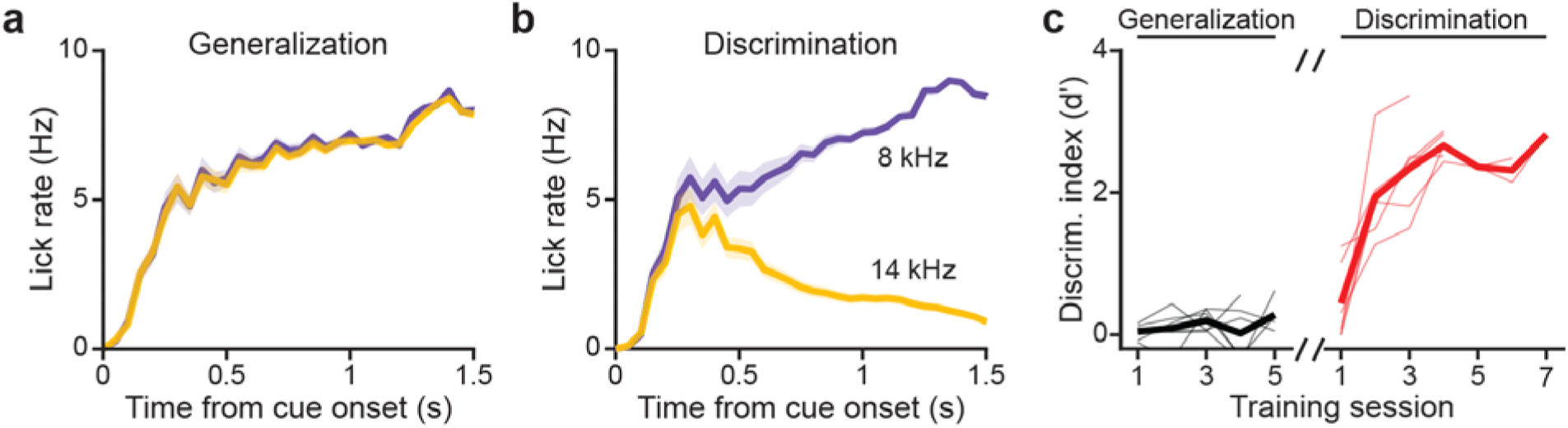
Additional data for behavioral performance during generalization and discrimination. **a**, **b**, Average lick rate for 8 and 14 kHz trials during imaging sessions in the generalization (**a**) and discrimination (**b**) stages. **c**, Discrimination performance (quantified as d’) across training sessions during generalization and discrimination. All error bands represent the s.e.m. and their centers represent the mean. *n* = 31 FOVs and 6 mice.

**Extended Data Fig. 2.**
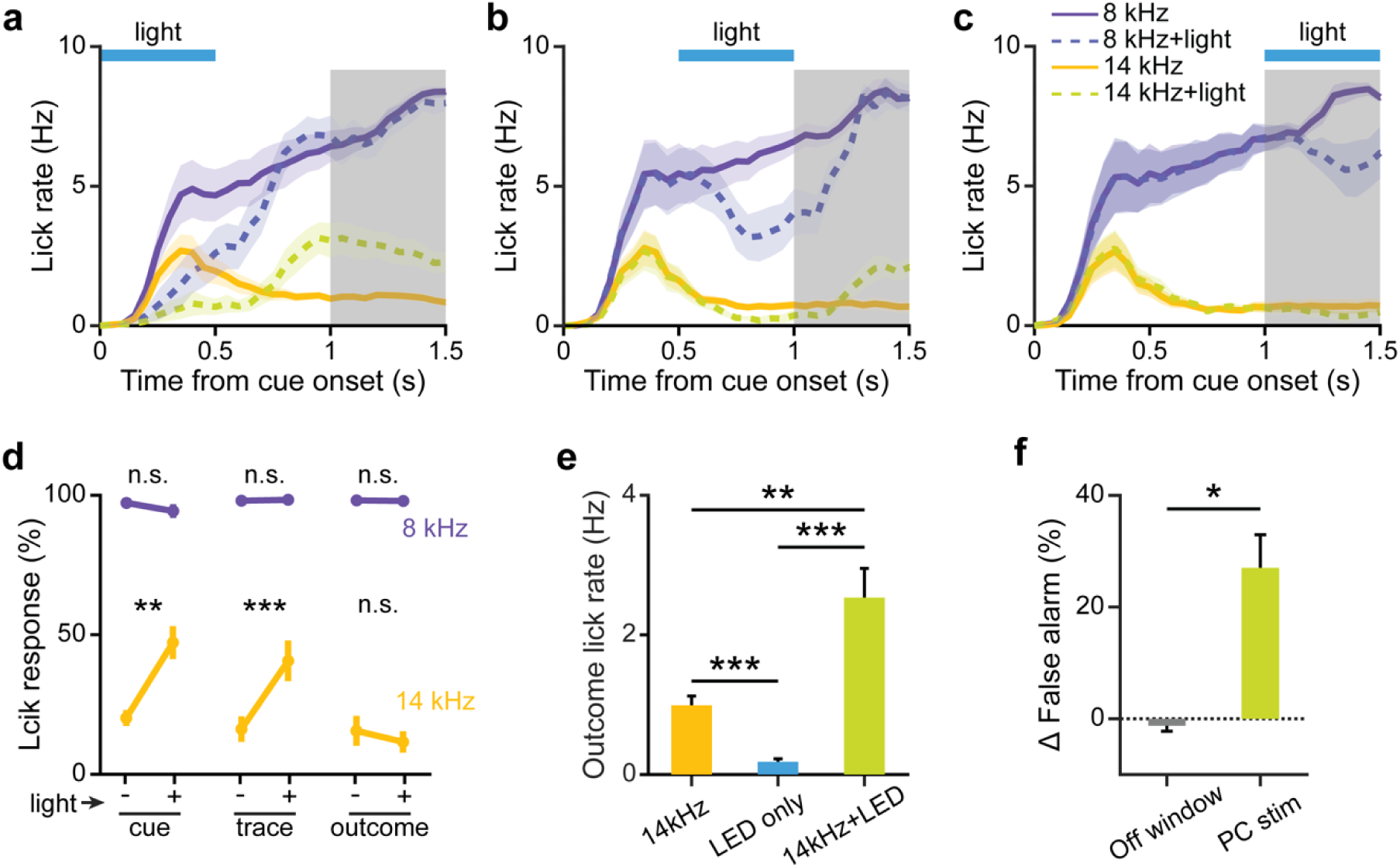
Additional data for optogenetic stimulation of Purkinje cells. **a**-**c**, Average lick rate for 8 and 14 kHz trials with and without PC stimulation during cue period (**a**), trace period (**b**), and outcome period (**c**). **d**, Percentage of trials with a lick response in outcome period (shaded in gray in **a**-**c**) for 8 kHz (Paired t-test, *P* = 0.070, *P* = 0.78, and *P* = 0.72, respectively) and 14 kHz (Paired t-test, *P* = 0.0026, *P* = 7.65 × 10^−4^, and *P* = 0.071, respectively) trials with or without PC stimulation during cue, trace, and outcome period. **e**, Lick rate during the outcome period in 14 kHz trials, LED-only control trials, and 14 kHz trials with PC stimulation. (Two sample t-test: 14 kHz vs LED-only, *P* = 4.34 × 10^−4^; LED-only vs 14 kHz + LED, *P* = 7.11 × 10^−4^; 14 kHz vs 14kHz + LED, *P* =0.0052) **f**, Difference in false alarm rates during regular PC stimulation and control experiments with LED fiber positioned outside the cranial window (Two sample t-test; *P* = 0.020). All error bars and error bands represent the s.e.m. and their centers represent the mean. n.s.: *P* > 0.05; \**P* ≤ 0.05; \*\**P* ≤ 0.01; \*\*\**P* ≤ 0.001. *n* = 8 mice.

**Extended Data Fig. 3.**
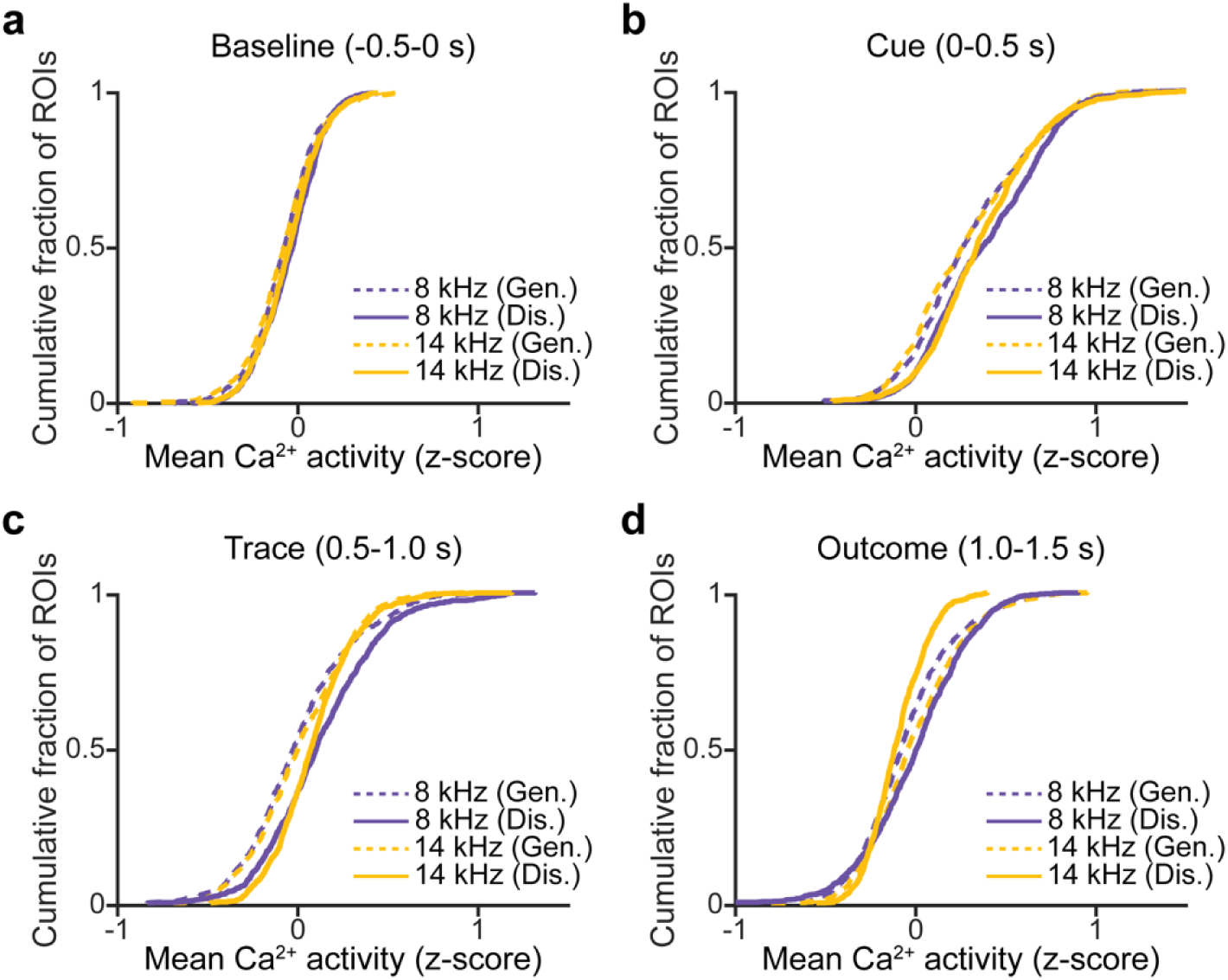
Supporting data for enhanced cue-evoked climbing fiber responses during sensory discrimination. **a-d**, Cumulative distribution frequency (CDF) plots of the mean calcium activity of all dendritic ROIs during baseline (**a**), cue (**b**), trace (**c**), and outcome (**d**) periods in 8 and 14 kHz trials across generalization and discrimination. *n* = 1010 ROIs for generalization and *n* = 1056 ROIs for discrimination.

**Extended Data Fig. 4.**
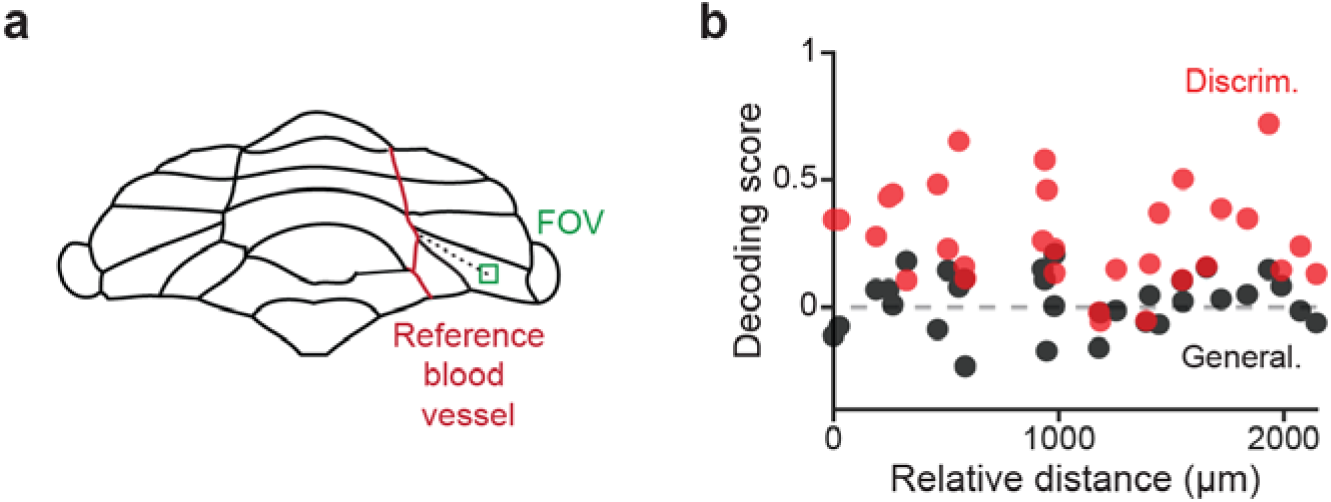
Tone decoding scores across FOV locations. **a**, Schematic illustrating the measurement of FOV distance relative to the reference blood vessel in the cerebellum. **b**, Correlation between decoding scores and FOV locations during generalization and discrimination. *n* = 31 FOVs.

**Extended Data Fig. 5.**
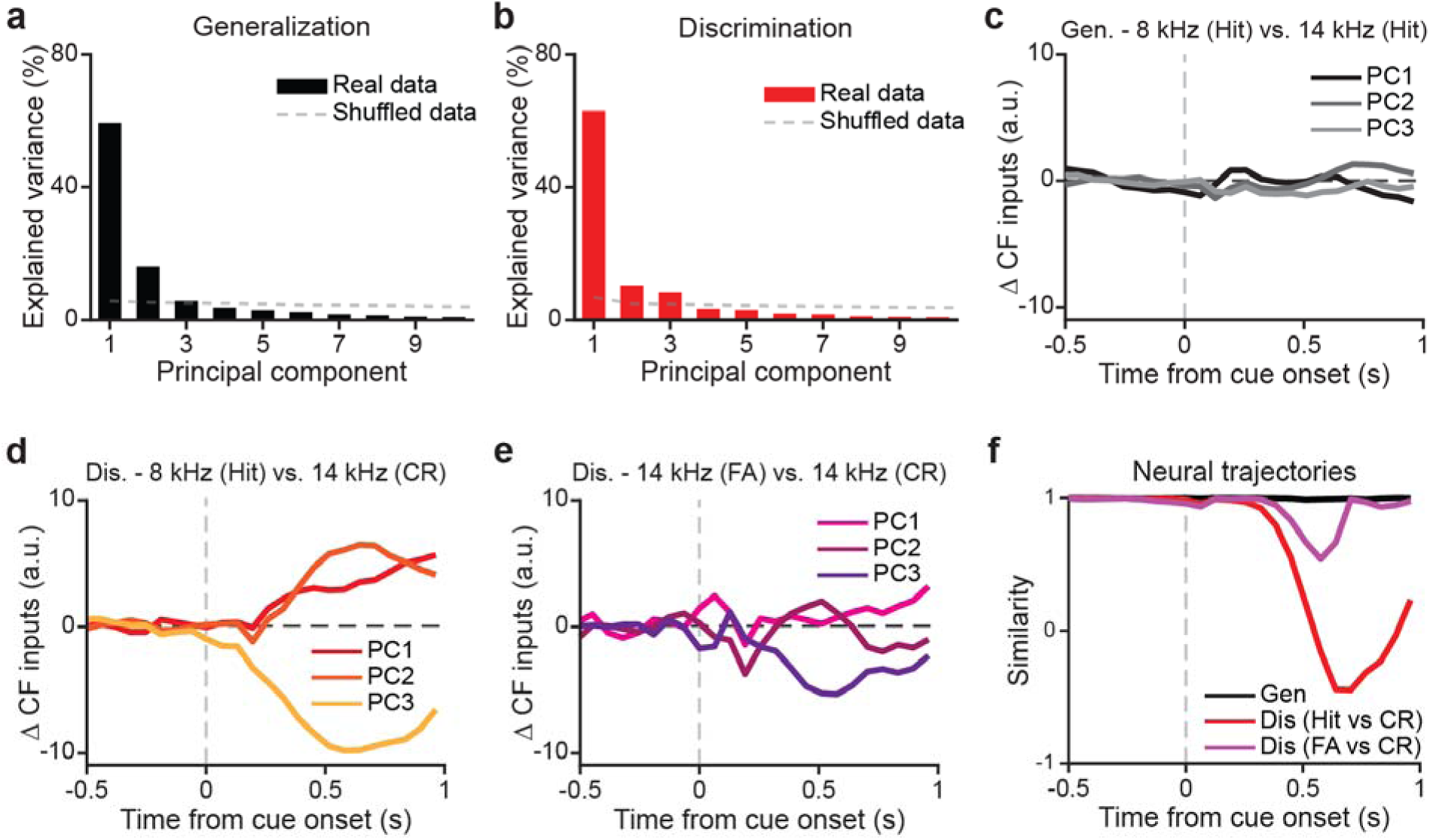
Additional data for decision-related neural trajectories analysis. **a**, **b**, **V**ariance explained by the top 10 principal components during generalization (**a**) and discrimination (**b**). **c**, Estimated differences in climbing fiber inputs for the first three principal components between correct 8 kHz and correct 14 kHz trials during generalization. **d**, Estimated differences in climbing fiber inputs for the first three principal components between hit and correct rejection trials during discrimination. **e**, Estimated difference in climbing fiber inputs for the first three principal components between false alarm and correct rejection trials during discrimination. **f**, Similarity of population neural activity trajectories for the comparisons in (**c-e**).

**Extended Data Fig. 6.**
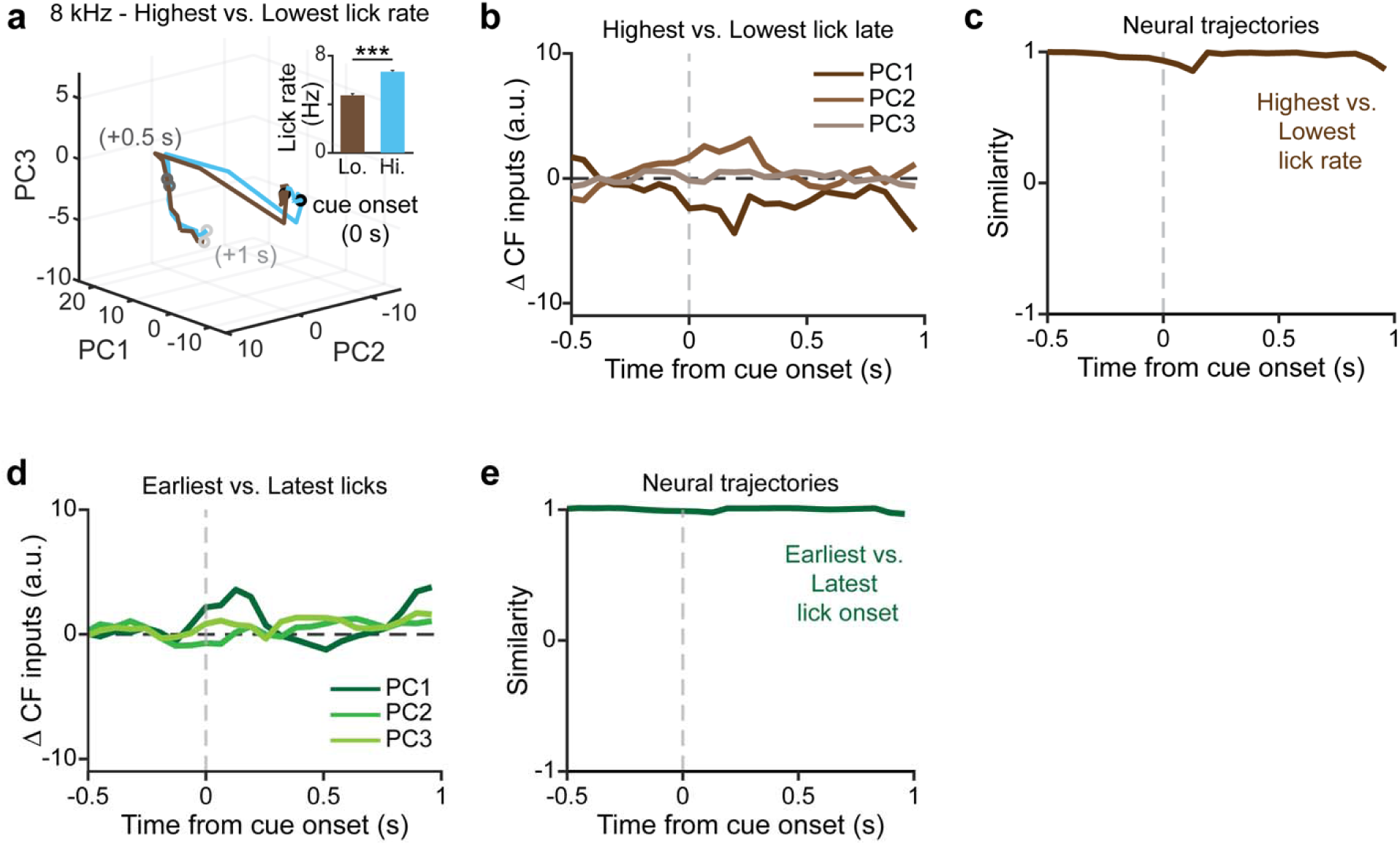
Additional data for lick-related neural trajectories analysis. **a**, Population neural activity trajectories for the lowest 25% and highest 25% lick rate 8 kHz trials during discrimination. Average lick rates are shown in inset (*n* = 280 and 311, respectively; Two sample t-test*, P* = 1.38 × 10^−25^). **b**, Estimated difference in climbing fiber inputs for the first three principal components between highest and lowest lick rate 8 kHz trials during discrimination. **c**, Similarity of population neural activity trajectories for the comparisons in (**b**). **d**, Estimated difference in climbing fiber inputs for the first three principal components between the earliest and the latest lick onsets 8 kHz trials during discrimination. **e**, Similarity of population neural activity trajectories for the comparisons in (**d**).

**Extended Data Fig. 7.**
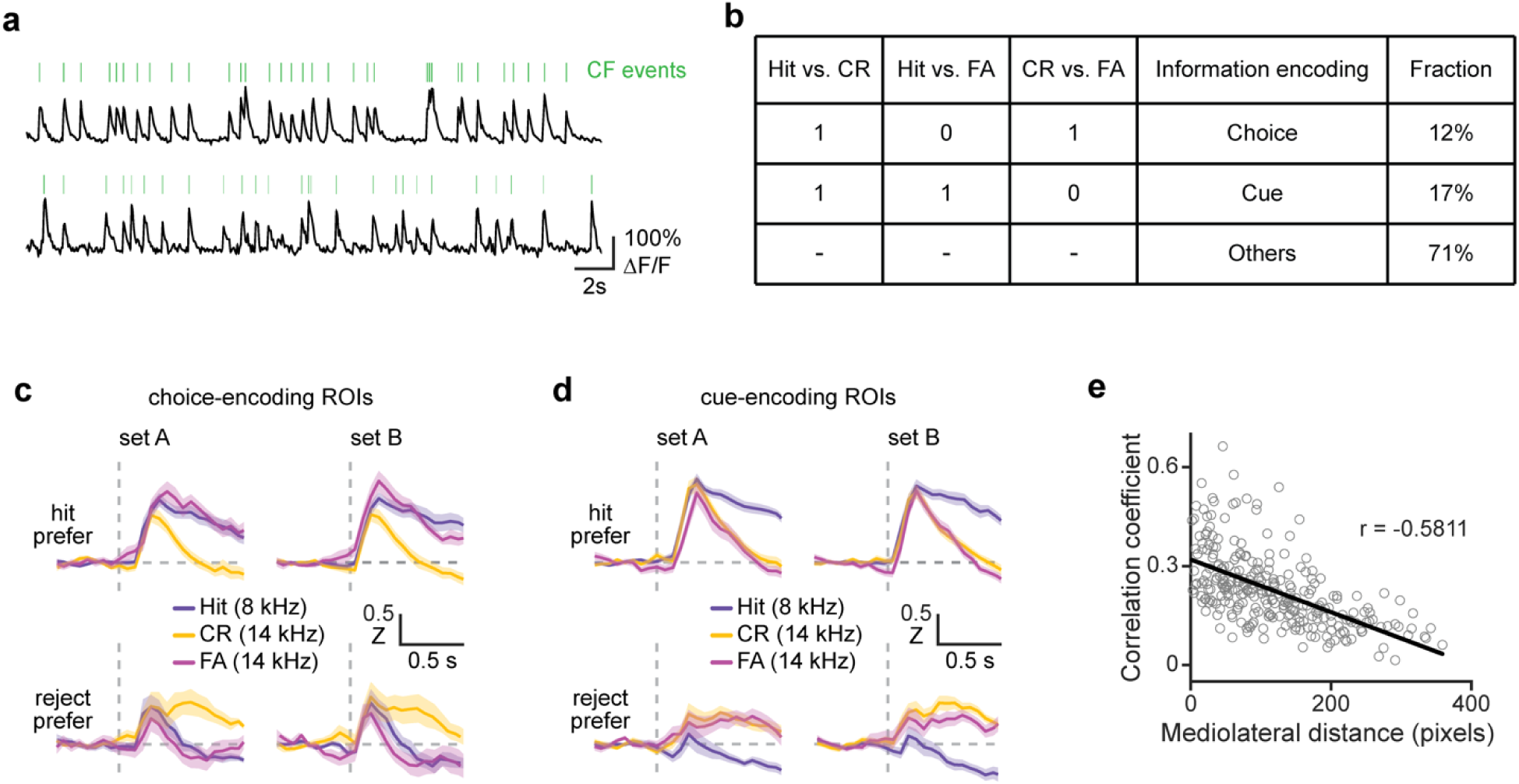
Additional data for information encoding properties of discriminative ROIs. **a**, Example fluorescence traces (black) and extracted calcium events (green) in two dendritic ROIs. **b**, Classification scheme of discriminative ROIs as either choice- or cue-encoding type. **c, d,** Trial-averaged traces of choice-encoding (**c**) and cue-encoding (**d**) ROIs from two randomly segregated subsets of trials. **e**, Correlation coefficient among dendritic ROI pairs as a function of mediolateral distance within an example FOV.

**Extended Data Fig. 8.**
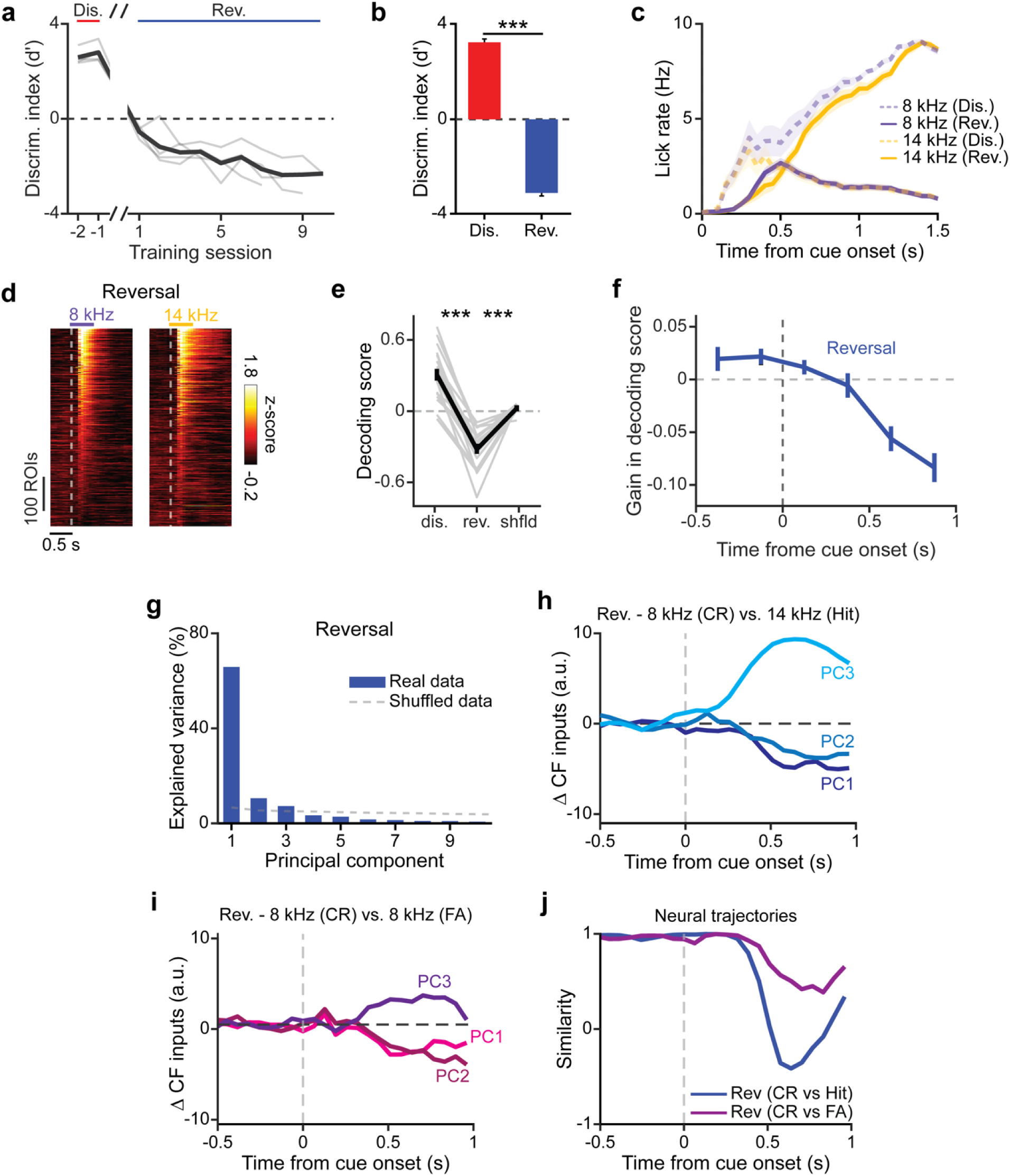
Additional data for reversal learning experiment. **a**, Discrimination performance (quantified as d’) across the last two training sessions of discrimination and all sessions of reversal. **b**, Behavioral performance (quantified as discrimination index d’) for all imaging sessions during discrimination and reversal. The same set of fields of view (FOVs) were examined (*n* = 19 from 4 mice; Two sample t-test; *P* = 6.05 × 10^−29^). **c**, Average lick rate during 8 and 14 kHz trials across discrimination and reversal imaging sessions (*n* = 19 from 4 mice). **d**, Raster plots of average calcium activity during 8 kHz and 14 kHz trials for all dendritic ROIs in generalization (left; *n* = 623) and reversal (right; *n* = 646). **e**, Changes in decoding scores for FOVs across discrimination and reversal, with tone labels shuffled as controls (*n* = 19, paired t-test, *P* = 1.96 × 10^−10^ and *P* =1.77 × 10^−7^, respectively). **f**, Temporal contributions of climbing fiber inputs to gain in decoding scores during reversal. **g**, **V**ariance explained by the top 10 principal components during reversal. **h**, Estimated difference in climbing fiber inputs for the first three principal components between correct rejection and hit trials during reversal. **i**, Estimated difference in climbing fiber inputs for the first three principal components between correct rejection and false alarm trials during reversal. **j**, Similarity of population neural activity trajectories for the comparisons in (**h)** and (**i)**.

**Extended Data Fig. 9.**
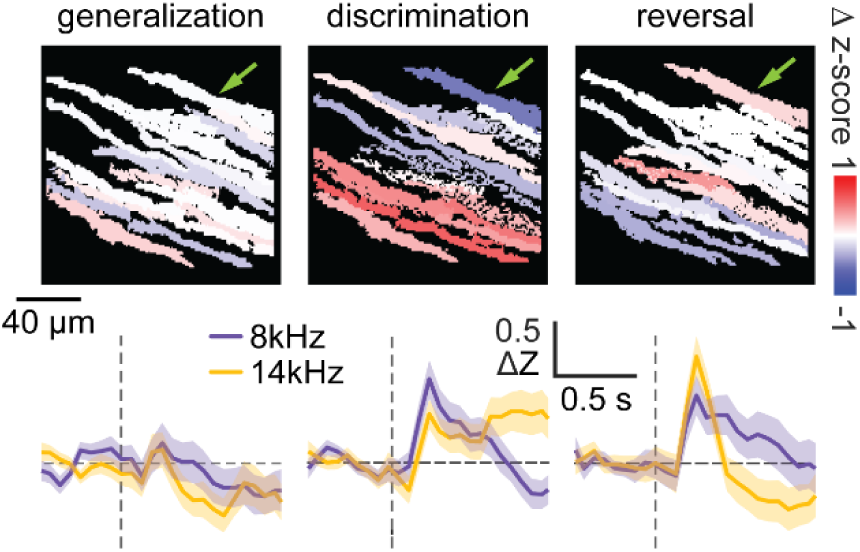
Additional data for plasticity in climbing fiber inputs of single Purkinje cell dendrite. Calcium response differences for matched ROIs in an example FOV (top) across stages and the trial-averaged traces (bottom) of an example polarity-switching ROI (green arrow).

**Extended Data Fig. 10.**
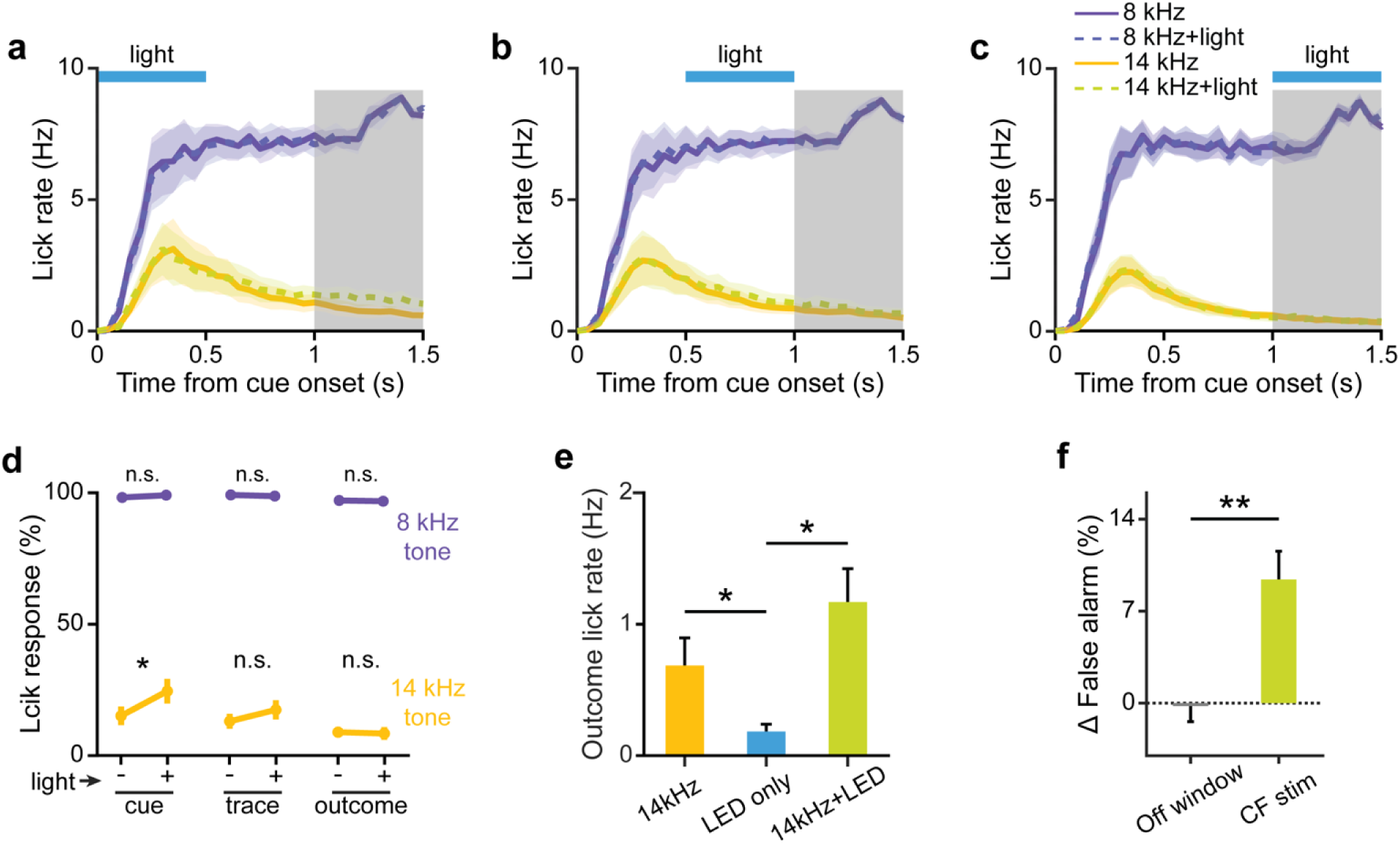
Additional data for optogenetic stimulation of climbing fibers. **a**-**c**, Average lick rate for 8 and 14 kHz trials with and without climbing fiber stimulation during cue period (**a**), trace period (**b**), and outcome period (**c**). **d**, Percentage of trials with a lick response in outcome period (shaded in gray in **a**-**c**) for 8 kHz (Paired t-test, *P* = 0.36, *P* = 0.57, and *P* = 0.69, respectively) and 14 kHz (Paired t-test, *P* = 0.0033, *P* = 0.12, and *P* = 0.71, respectively) trials with or without climbing fiber stimulation during cue, trace, and outcome period. **e**, Lick rate during the outcome period in 14 kHz trials, LED-only control trials, and 14 kHz trials with climbing fiber stimulation (Two sample t-test: 14 kHz vs LED-only, *P* = 0.048; LED-only vs 14 kHz + LED, *P* = 0.0033; 14 kHz vs 14kHz + LED, *P* = 0.1922). **f**, Difference in false alarm rates during regular climbing fiber stimulation and control experiments with LED fiber positioned outside the cranial window (Two sample t-test; *P* = 0.0015). All error bars represent the s.e.m. and their centers represent the mean. n.s.: *P* > 0.05; \**P* ≤ 0.05, \*\**P* ≤ 0.01. *n* = 8 mice.

## Notes

### Competing Interest Statement

The authors have declared no competing interest.

## References

1. Kim, S.-G., Uğurbil, K. & Strick, P. L. Activation of a Cerebellar Output Nucleus During Cognitive Processing. Science (80-. ). 265, (1994).

2. Gao, J. H. et al. Cerebellum implicated in sensory acquisition and discrimination rather than motor control. Science (80-. ). 272, (1996).

3. Schmahmann, J. D. The cerebellum and cognition. Neuroscience Letters vol. 688 at 10.1016/j.neulet.2018.07.005 (2019).

4. Gilbert, P. F. C. & Thach, W. T. Purkinje cell activity during motor learning. Brain Res. 128, (1977).

5. Ito, M. & Kano, M. Long-lasting depression of parallel fiber-Purkinje cell transmission induced by conjunctive stimulation of parallel fibers and climbing fibers in the cerebellar cortex. Neurosci. Lett. 33, (1982).

6. Schultz, W. Behavioral theories and the neurophysiology of reward. Annu. Rev. Psychol. 57, 87–115 (2006).

7. Heffley, W. et al. Coordinated cerebellar climbing fiber activity signals learned sensorimotor predictions. Nat. Neurosci. 21, (2018).

8. Kostadinov, D., Beau, M., Pozo, M. B. & Häusser, M. Predictive and reactive reward signals conveyed by climbing fiber inputs to cerebellar Purkinje cells. Nat. Neurosci. 22, (2019).

9. Heffley, W. & Hull, C. Classical conditioning drives learned reward prediction signals in climbing fibers across the lateral cerebellum. Elife 8, (2019).

10. Ishikawa, T., Shimuta, M. & Häuser, M. Multimodal sensory integration in single cerebellar granule cells in vivo. Elife 4, (2015).

11. Ju, C. et al. Neurons of the inferior olive respond to broad classes of sensory input while subject to homeostatic control. J. Physiol. 597, (2019).

12. Ozden, I., Sullivan, M. R., Lee, H. M. & Wang, S. S. H. Reliable coding emerges from coactivation of climbing fibers in microbands of cerebellar Purkinje neurons. J. Neurosci. 29, (2009).

13. Kitamura, K. & Häusser, M. Dendritic calcium signaling triggered by spontaneous and sensory-evoked climbing fiber input to cerebellar purkinje cells in vivo. J. Neurosci. 31, (2011).

14. Pachitariu, M., et al. Suite2p: beyond 10,000 neurons with standard two-photon microscopy. bioRxiv (2016) doi:10.1101/061507.

15. Koren, V., Andrei, A. R., Hu, M., Dragoi, V. & Obermayer, K. Pairwise Synchrony and Correlations Depend on the Structure of the Population Code in Visual Cortex. Cell Rep. 33, (2020).

16. Geiller, T. et al. Large-Scale 3D Two-Photon Imaging of Molecularly Identified CA1 Interneuron Dynamics in Behaving Mice. Neuron 108, (2020).

17. Deneux, T. et al. Accurate spike estimation from noisy calcium signals for ultrafast three-dimensional imaging of large neuronal populations in vivo. Nat. Commun. 7, (2016).

18. Apps, R. & Garwicz, M. Anatomical and physiological foundations of cerebellar information processing. Nature Reviews Neuroscience vol. 6 at 10.1038/nrn1646 (2005).

19. Bonnan, A., Rowan, M. M. J., Baker, C. A., Bolton, M. M. L. & Christie, J. M. Autonomous Purkinje cell activation instructs bidirectional motor learning through evoked dendritic calcium signaling. Nat. Commun. 12, (2021).

20. Wolpert, D. M., Miall, R. C. & Kawato, M. Internal models in the cerebellum. Trends in Cognitive Sciences vol. 2 at 10.1016/S1364-6613(98)01221-2 (1998).

21. Streng, M. L., Popa, L. S. & Ebner, T. J. Climbing fibers predict movement kinematics and performance errors. J. Neurophysiol. 118, (2017).

22. Pisano, T. J. et al. Homologous organization of cerebellar pathways to sensory, motor, and associative forebrain. Cell Rep. 36, (2021).

23. Oscarsson, O. Functional units of the cerebellum - sagittal zones and microzones. Trends in Neurosciences vol. 2 at 10.1016/0166-2236(79)90057-2 (1979).

